# Relationship between monomer packing, receptor binding domain pocket status, and pH, in the spike trimer of SARS-CoV-2 variants

**DOI:** 10.1101/2021.12.14.472554

**Authors:** Jim Warwicker

## Abstract

Existence of a SARS-CoV-2 spike protein trimer form with closer packing between monomers when receptor binding domains (RBDs) are all down, locked as opposed to closed, has been associated with linoleic acid (LA) binding at neutral pH, or can occur at acidic pH in the absence of LA binding. The relationship between degree of closure of the LA binding pocket of the RBD, and monomer burial in the trimer, is examined for a range of spike protein structures, including those with D614G mutation, and that of the Delta variant (which also carries D614G). Some spike protein structures with this aspartic acid mutation show monomer packing approaching that of the locked form (at neutral pH, without LA binding) for two segments, a third (around the RBD) remains less closely packed. Mutations in the RBD are a focus for the Omicron variant spike protein. Structure reports suggest that these mutations are involved in increased RBD-RBD interactions, and also that they could lead to a closing of the LA pocket, both of which could impact on pH-dependence. One potential outcome is that the extent of pH-dependent conformational transitions of the pre-fusion SARS-CoV-2 spike trimer are reduced in the Omicron variant.

## Introduction

Viruses must enter a cell and release their genome for copying, they must also exit the cell. For both entry and exit, pH-dependent processes may play a role, including receptor-mediated endocytosis into low pH endosomes and navigation of acidic pH in the secretory pathway (Robinson et al., 2018). For the membrane enveloped coronavirus family, cell entry may be through fusion directly at the cell membrane or through fusion at an intracellular membrane, subsequent to receptor-mediated endocytosis of the virus (Whittaker et al., 2021). Although there is evidence that altering endosomal pH impedes viral entry to some extent (Prabhakara et al., 2021) for the SARS-CoV-2 coronavirus (causative agent of the covid-19 pandemic), the precise balance of genome release routes (cell surface or interior) may depend on other factors, such as the priming cleavage of S1 and S2 subunits of the spike (S) protein (Papa et al., 2021). For membrane enveloped viruses, mechanisms evolve to protect against mis-timed low pH-induced membrane fusion events in the acidic pH secretory pathways for newly synthesised virus (Fields and Kielian, 2015). It is apparent that questions of pH-dependence in these systems are complex, relate to pH inside and outside of the cell, and couple to other determinants of stability, including receptor and other ligand binding, and proteolytic cleavages. A strong signal in calculations of pH-dependence, relative to the unfolded state, for each of the pre and post-fusion structures of SARS-CoV-2 S protein, are three His sidechains in the S2 subunit, but since their burial and predicted pH-dependence are uniform between pre and post-fusion forms, it is unclear whether they are functionally relevant (Warwicker, 2021). For coronaviruses, conformational variation of pre-fusion S protein is apparent from cryo-EM structures. Indeed, there are major questions to be answered about the pH-dependence of S1/S2 subunit S-protein structure and function, even without considering the large-scale conformational changes of the S2 subunit that follow S1 subunit release, and accompany membrane fusion (Cai et al., 2020).

Based on the response of S protein trimer structure to acidic pH (Zhou et al., 2020; Qu et al., 2021), a scheme has been suggested in which the trimers on a virus within the acidic pH of the secretory pathway would be protected in a more tightly packed (locked) conformation (Qu et al., 2021). This suggestion is consistent with the discovery of a locked and more tightly packed form of SARS-CoV-2 S protein trimer at the normal pH (7.5 to 8) of structure solution, due to binding of a linoleic acid pocket factor in the receptor binding domain (RBD) (Toelzer et al., 2020). These observations prompt questions of which S protein amino acids are sensing pH, and how these might be coupled to pocket factor binding and packing/interface changes. A study of S protein trimers grouped according to RBD down (with locked and closed forms treated separately), and RBD up, reported differences in the predicted pH-dependence (Lobo and Warwicker, 2021). The largest contributions to these differences arose from interface tightening in the trimer between closed and locked forms, rather than the well-known difference between open (RBD up) and closed (RBD down) forms, and specifically from Asp and Glu sidechains, prominent amongst which is D614. It was proposed that these carboxylate groups have relatively unperturbed p*K*_*a*_s in open and closed forms, but that increased burial in locked forms leads to destabilisation. Depending on the balance of overall interface energetics, the effect could be to favour locked forms at acidic pH (less destabilised carboxylates, pH closer to their normal p*K*_*a*_), and closed/open forms at neutral pH (since locked form carboxylates would be more destabilised, with pH further from their normal p*K*_*a*_) (Lobo and Warwicker, 2021), a similar proposal to that of protection in the secretory pathway (Qu et al., 2021).

The picture for SARS-CoV-2 spike protein encountered in the first year of the covid-19 pandemic is therefore of a trimer that can exist in a more tightly packed form (termed locked) either with pocket factor binding at neutral pH, or at acidic pH in the absence of pocket factor. However, the spike protein is changing, leading to the question of what the consequences may be for relative stability of the locked, closed, and open forms.

## Materials and Methods

Inspection of the spike protein LA binding pocket (Fig. 1) revealed that its open/closed state could be characterised by a distance between the average of the 4 β-strand C_α_ atoms on one side of the pocket (amino acids K378, C379 and C432, V433), and the average of the two C_α_ atoms on the opposite side (amino acids L368, Y369). Whereas the 4 β-strand amino acids on one side, including a disulphide bridge between C379 and C432), are structurally relatively invariant, L368, Y369 are in a turn that is part of the gating mechanism for the pocket (Toelzer et al., 2020).

**Figure 1.**
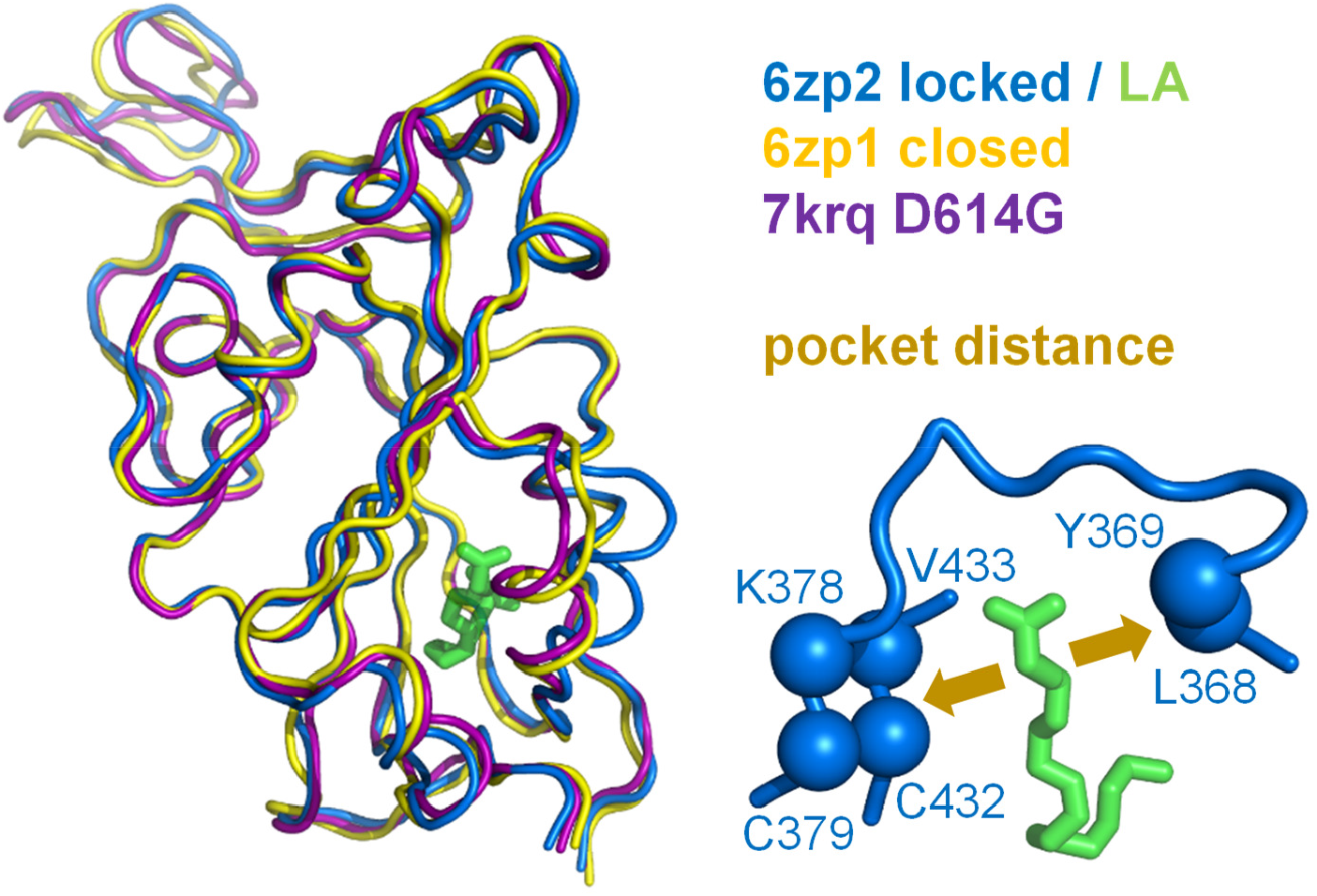
A measure of pocket opening in the RBD. Receptor binding domains are aligned for representative locked (6zp2) and closed (6zp1) (Xiong et al., 2020) S protein trimers, and also for an S trimer structure carrying the D614G mutant, 7krq (Zhang et al., 2021a), with all RBDs down and monomer burial approaching the locked form. Distance across the pocket is shown schematically for 6zp2, with linoleic acid (LA) bound. The distance is calculated between the average of the 4 β-strand C_α_ atoms displayed (left of LA, which align well structurally between these RBDs), and the average of the two C_α_ atoms shown to the right of LA, present on a turn within a sub-structure that gates the LA binding pocket (Toelzer et al., 2020).

Burial of spike protein monomer within a trimer for structures of Delta variant S protein was calculated as described previously for other SARS-CoV-2 variant trimers (Lobo and Warwicker, 2021), with d-SASA equal to the change in solvent accessible surface area between free monomer and monomer within a trimer. These d-SASA values are used either as averages over monomers within one trimer structure, or as averages over all monomers within a specified sub-grouping of structures. When plotted as profiles over the spike protein sequence, a further difference is made, between d-SASA profiles, to give a direct indication of where packing varies between systems.

RBD structures for other coronaviruses were examined by searching the SARS-CoV-2 spike sequence, id P0DTC2 from UniProt (UniProt, 2019) against the RCSB/PDB (Berman et al., 2007) with the Basic Local Alignment Search Tool (BLAST) (Altschul et al., 1990) at the National Center for Biotechnology Information. Protein structures were aligned with Swiss PDB Viewer (Guex and Peitsch, 1997) and visualised with Swiss PDB Viewer and PyMol.

## Results and Discussion

Previous predictions of potential acid sensor residues, and calculations of the change in solvent accessible surface area (d-SASA) upon spike monomer incorporation in trimer (in various trimer forms) (Lobo and Warwicker, 2021), are supplemented with data for two recent structures of the Delta variant (RCSB/PDB (Berman et al., 2007) identifiers 7v7n (10.2210/pdb7V7N/pdb) and 7sbk (Zhang et al., 2021b)), and consideration of sequence changes around the RBD pocket of the Omicron variant S protein. A new measure is constructed of distance across the RBD pocket in which linoleic acid can bind (Fig. 1). Comparing pocket distance and monomer interface in a scatter plot (Fig. 2), it is apparent that open and closed forms cluster with a distance consistent with a closed pocket, and at low (open) and intermediate (closed) monomer burial in the trimer. Locked forms are quite separate, with a larger (and occupied) pocket, and high monomer burial. The set of pH-locked forms have closed pockets, and monomer burial intermediate between closed and locked forms, but at acidic pH (all other data in the plot is for structures at or slightly above neutral pH). Structures in both the D614Gset-closed and Delta-closed groups carry the D614G mutation, and have been filtered for trimers in which all RBDs are in the down conformation. For both groups, the clustering exhibited by other forms is lost, monomer burial in the trimer can vary substantially, and one structure also has an intermediate pocket distance. Variants with the S protein D614G mutation increase recovery in cryo-EM structure determination of forms that approach a locked degree of monomer interface burial, but at neutral rather than acidic pH, with evidence that they are also able to sample conformations of intermediate pocket closure (Fig. 2). It seems that the barrier between locked and closed forms at neutral pH is reduced by the D614G substitution.

**Figure 2.**
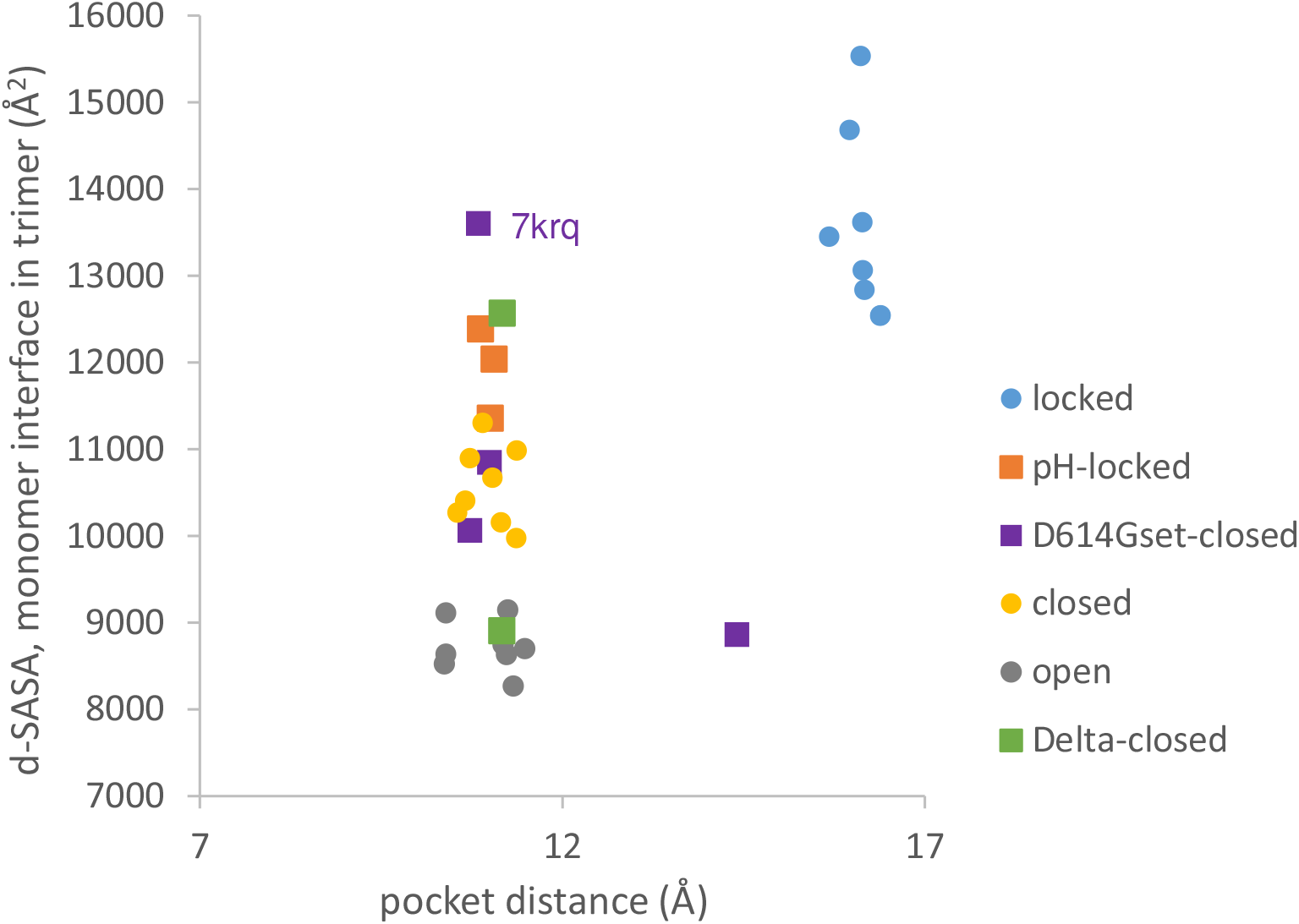
Variants carrying the D614G mutation do not cluster on a plot of pocket distance against S monomer burial within the S trimer. Clustering of locked, pH-locked, closed, and open forms in terms of monomer burial (Lobo and Warwicker, 2021) extends also to pocket distance. However, D614G set S proteins show greater variation, and the two Delta variant S proteins displayed are also well separated in monomer burial. Values are 3 monomer averages for each of the S protein trimers. Other than the open set, only trimers with all RBDs down are included (and thus sets are named D614Gset-closed and Delta-closed).

In order to establish the location of interface differences (for monomer burial in trimer), and how they change in a D614G structure with similar burial to locked forms, differences between 7krq (Zhang et al., 2021a) and (averaged) locked form burial are studied (Fig. 3). There is an interaction of equivalent helices in the trimer towards the C-terminus, which is only present for some structures. Other than this, major regions of interaction differences between 7krq and the closed form (555 – 670) and (830 – 860) have, by contrast, similar monomer burial in 7krq (neutral pH, D614G mutation, pocket closed) and the locked form (neutral pH, linoleic acid bound). These two regions contact between monomers in the trimer, and it has been suggested that a salt-bridge lost between D614 and K854 (of neighbouring monomers) in variants carrying the D614G mutation, effectively breaks a latch and leads to a greater fraction of open (RBD up) forms, thereby enhancing interaction with the receptor (Yurkovetskiy et al., 2020). Another interpretation of the effect of D614G mutation is that it stabilises the S protein trimer against dissociation (Zhang et al., 2021a), which is consistent with the hypothesis that once a destabilising burial of D614 is lost, a locked form is more accessible at neutral pH (Lobo and Warwicker, 2021).

**Figure 3.**
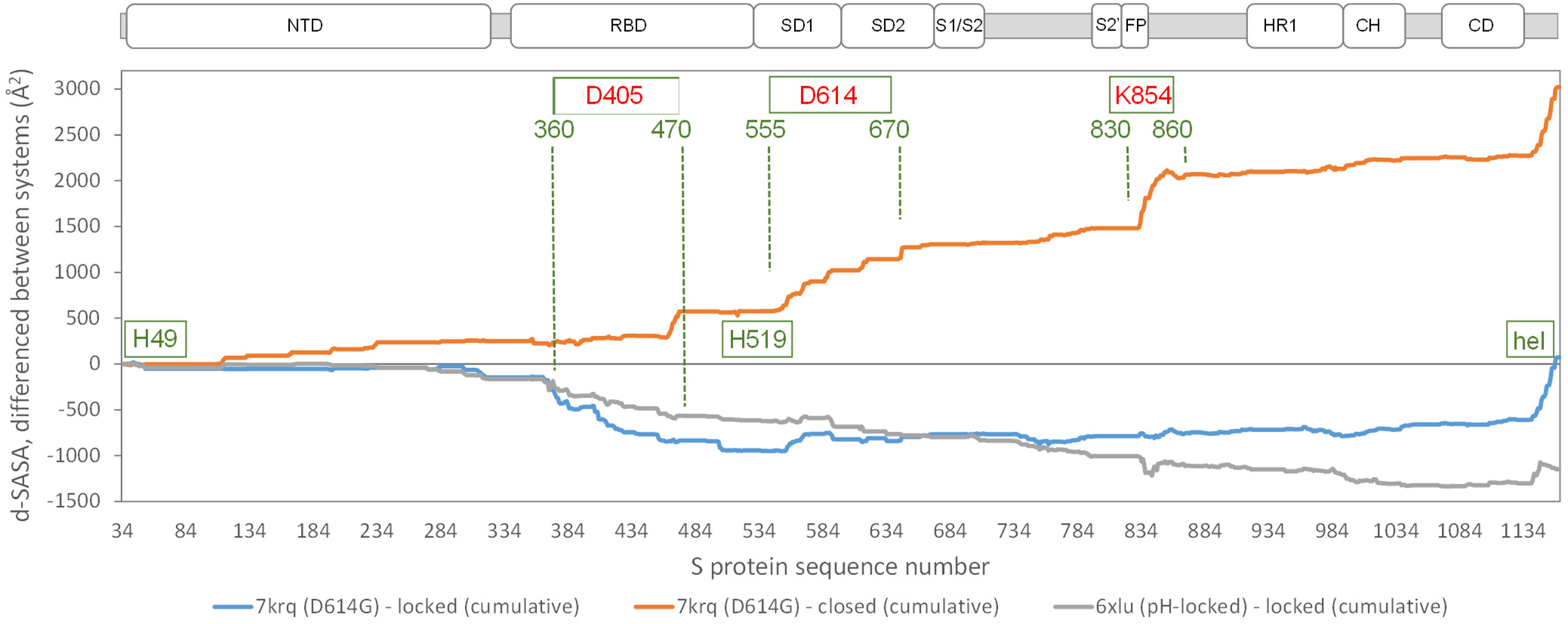
Comparison of acidic pH and D614G mutation effects on packing within the S trimer. The d-SASA value (monomer burial within a trimer) is further differenced between the systems indicated, between 7krq (D614G, closed pocket but burial approaching the locked form) and the average over the locked set, between 7krq and the average over the closed set, and between 6xlu (pH-locked) and the average over the locked set. These double difference quantities are plotted cumulatively over the sequence of the S trimer, for which structure is available, with sub-domains (Berger and Schaffitzel, 2020) indicated (NTD N-terminal domain, SD1/SD2 subdomains 1 and 2, S1/S2 proteolytic cleavage site between subunits, S2’ cleavage site within S2 subunit, FP fusion peptide, HR1 heptad repeat 1, CH central helix, CD connector domain). Regions and amino acids of particular interest are displayed with residue numbers and in relation to the changes in burial. A C-terminal helical coil interaction between monomers (present only in some structures) is labelled (hel).

A further region of interest is (360 – 470), where there is most differentiation between the D614G 7krq structure and the locked / pocket occupied form (Fig. 3). This segment is of interest for several reasons. Within it lies parts of the linoleic acid pocket, including the gating mechanism (Toelzer et al., 2020) (Fig. 1). It also contains carboxylate groups of D405, D420, and E465 that are proposed (together with D614) to couple burial and pH-dependent stability (Lobo and Warwicker, 2021). Interestingly, in this region, the acid pH-locked structure 6xlu (Zhou et al., 2020) is more similar to the neutral pH locked form than is the D614G 7krq structure (Fig. 3). If there were one or more buried and destabilised Asp/Glu sidechains in this region, then such behaviour would be expected.

Finally, S protein of the Omicron variant (https://www.ecdc.europa.eu/en/covid-19/variants-concern) carries S371L, S373P, S375F mutations in the sequence next to the RBD pocket gate (Fig. 4). There are many further mutations in Omicron S protein, which are being discussed in the context of various features, including the extent to which they could mediate escape of an immune response primed by vaccine or prior infection (Greaney et al., 2021; Zahradník et al., 2021), and how immune escape mutations may be compensated to maintain ACE2-binding (Mannar et al., 2021) (Pengcheng et al., 2022). Interesting mutations in the Omicron spike protein, with respect to packing, include D614G and the S371L, S373P, S375F cluster. The position equivalent to 373 in the RBD of coronavirus HKU9 is also a proline, and since this amino acid lies in a segment connecting two sides of the LA-binding pocket (in SARS-CoV-2 spike protein), a proline substitution could alter mainchain conformation and pocket gating. Indeed, the equivalent pocket region in HKU9 (5gyq) (Huang et al., 2016) looks to be closed, with a similar outcome anticipated for the RBD of SARS-CoV-2 Omicron variant (Zahradník et al., 2021). If this pocket is closed in the Omicron variant spike, then linoleic acid would not bind and thus would not be a route to locking (compacting) the spike protein structure. It is intriguing that early reports of Omicron spike structure indicate a capacity to form relatively compact structures in both closed (all RBD down) and open forms (Hong et al., 2022). Reports suggest that for the 1 RBD up form this may be due, in part, to enhanced interactions between monomers (S375F – F486) (Zhou et al., 2021).

**Figure 4.**
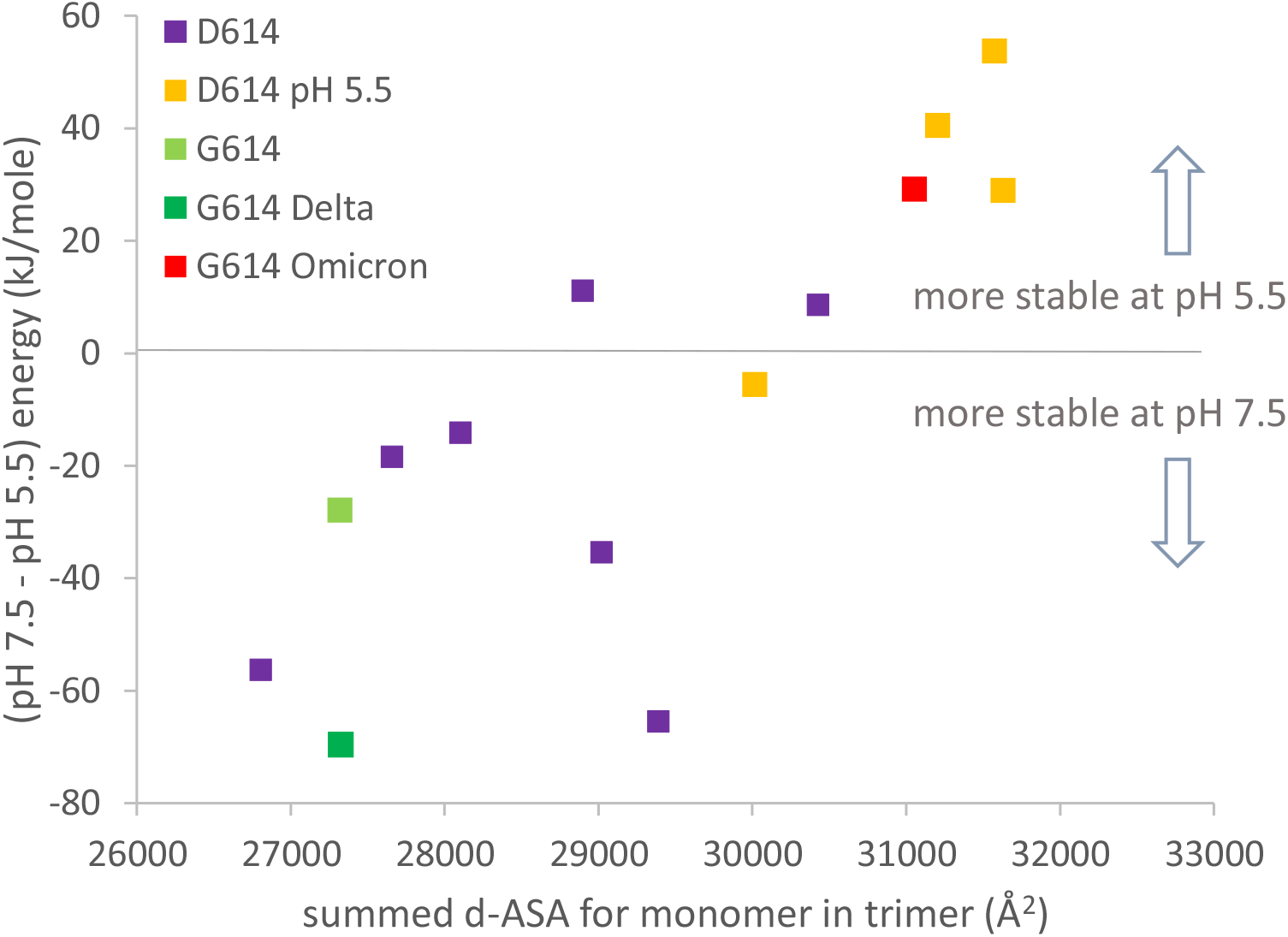
Compaction of the 1 RBD up trimer correlates with prediction of altered pH-dependent stability. Engineered (2P) 1 RBD up structures are (D614: 6xf6, 7knb, 7kne, 6xm3, 6xm4, 6xm0, 6vsb, 7byr, 6vyb, 7cn9, 6×2x; G614: 7bnn; G614-Delta 7w98; G614-Omicron 7tb4). d-ASA for each structure is summed over the 3 monomers in a trimer. Predicted pH-dependence of trimer structural stability is calculated, following reported methods (Lobo and Warwicker, 2021), between pH 7.5 and pH 5.5.

The major thrust of pH-dependence predictions for non-Omicron variants is that increased compaction of spike trimer leads to partial dehydration of some Asp/Glu sidechains, and a disfavouring of a more compact form at neutral pH, unless balanced by linoleic acid binding to the RBD pocket (Lobo and Warwicker, 2021). In order to include the Omicron variant spike protein (which may not bind linoleic acid) in this picture, it is necessary to study the 1 RBD up form, since this is the only currently available structure (January 2021) that has a complete set of 3 RBDs resolved (7tb4) (Zhou et al., 2021). Further, to maintain equivalence between engineered status of the spike protein ectodomain, analysis is restricted to the 2P (K986P, V987P) stabilized form. In keeping with previous work (Lobo and Warwicker, 2021), increased trimer compaction correlates with a relative lowering of stability at pH 7.5 compared with pH 5.5 (Fig. 4). The 1 RBD up Omicron spike protein is predicted to be as compact as 1 RBD up trimers that have been stabilized by lowering the pH of structure solution to 5.5, although the Omicron spike was not solved at acidic pH. Presumably this increased compaction arises from interactions (including S375F – F486) that are absent in other SARS-CoV-2 variant spike proteins, and that balance the predicted electrostatic destabilization at neutral pH. Since relatively compact trimers have been reported for both all RBD down and RBD up Omicron spike protein structures at neutral pH (Hong et al., 2022), likely in the absence of RBD pocket factor binding, it is reasonable to suggest that they may be better protected against S1 shedding than spike trimer of other SARS-CoV-2 variants. Further, although a predicted pH-dependence of stability remains for the Omicron variant spike, with an overall shift in the balance of trimer stabilizing interactions, it is unclear how this will translate to the prevalence of different spike conformational forms in different pH environments. One possibility is that with relatively compact closed and open (at least 1 RBD up) forms, with similar electrostatic destabilization (due to partial Asp/Glu burial), transition between the forms would be facilitated.

## Conclusion

Although structural studies of SARS-CoV-2 spike proteins continue at pace, many mechanisms remain unknown, including that behind the pH-dependence of S trimer conformation, and how it might relate to function. The proposal that Asp/Glu sidechains could be responsible for observed pH-dependent effects is yet to be tested. An alternate hypothesis is that histidines H49 and H519 could be mediating pH-dependence (Qu et al., 2021), but these give small predicted pH-dependence (Lobo and Warwicker, 2021) and are located in regions without large interface change between structural forms (Fig. 3). Here the degree to which the D614G mutation allows access of spike trimers (without LA binding) to conformations that are similar to the LA-bound locked conformations, is assessed. Analysis of mutations around the gate for the LA binding pocket (Toelzer et al., 2020), in the Omicron variant spike protein, suggests that further changes may occur, towards locked conformation accessibility at neutral pH (and without LA binding). The 1 RBD up open form of the Omicron spike trimer appears to be more compact at neutral pH than it is for other variants, in part due to altered interactions between RBDs (Zhou et al., 2021). It is possible that a hypothesised role for pH-dependence in regulating conformational transitions of the pre-fusion SARS-CoV-2 spike trimer (Lobo and Warwicker, 2021; Qu et al., 2021) is reduced in the Omicron variant, where both all RBD down and 1 RBD up forms are reported to be relatively compact at neutral pH (Hong et al., 2022).

## Acknowledgements

The author thanks staff who support the computational shared facility (CSF) at the University of Manchester.

